# Agent-based network model predicts strong benefits to youth-centered HIV treatment-as-prevention efforts

**DOI:** 10.1101/207126

**Authors:** John E Mittler, James T Murphy, Sarah Stansfield, Kathryn Peebles, Geoffrey S Gottlieb, Steven Goodreau, Joshua T Herbeck

## Abstract

We used an agent-based network model to examine the effect of targeting different risk groups with unsuppressed HIV viral load for linkage or re-linkage to HIV-related treatment services in a heterosexual population with annual testing. Our model identifies prevention strategies that can reduce incidence to negligible levels (i.e., less than 0.1 infections per 100 person-years) 20 years after a targeted Treatment-as-Prevention (TasP) campaign. The model assumes that most (default 95%) of the population is reachable (i.e., could, in principle, be linked to effective care) and a modest (default 5% per year) probability of a treated person dropping out of care. Under random allocation or CD4-based targeting, the default version of our model predicts that the TasP campaign would need to suppress viral replication in ~80% of infected people to halt the epidemic. Under age-based strategies, by contrast, this percentage drops to 50% to 60% (for strategies targeting those <30 and <25, respectively). Age-based targeting did not need to be highly exclusive to yield significant benefits; e.g. the scenario that targeted those <25 years old saw ~80% of suppressed individuals fall outside the target group. This advantage to youth-based targeting remained in sensitivity analyses in which key age-related risk factors were eliminated one by one. As testing rates increase in response to UNAIDS 90-90-90 goals, we suggest that efforts to link all young people to effective care could be an effective long-term method for ending the HIV epidemic. Linking greater numbers of young people to effective care will be critical for developing countries in which a demographic “youth bulge” is starting to increase the number of young people at risk for HIV infection.

---

UNAIDS and other public health agencies have promoted the 90-90-90 targets for HIV diagnosis, treatment, and viral suppression by 2020 (UNAIDS 2017). However, the resulting ~73% viral suppression may not be enough to end the epidemic; in regions with high *R*_0_’s (i.e., regions where the virus can spread quickly) mathematical models predict that 80% to 90% of HIV-infected individuals will need to be virally suppressed to reduce incidence to zero (Kretzschmar *et al.* 2013, Williams and Gouws 2013, Hontelez *et al.* 2013, Rozhnova *et al.* 2016). For populations with very high *R*_0_ values (Williams and Gouws 2013, Kretzschmar *et al.* 2013, Rozhnova *et al.* 2016), even a more ambitious “95-95-95 target” (~86% viral suppression) (Medlock *et al.* 2017) may not be enough. Alsallaq et al (2017) note, however,that “youth-focused” prevention and treatment programs can outperform “adult-focused” programs, suggesting a more effective approach for HIV eradication. While age-based HIV testing has been modeled (Bershteyn *et al.* 2016, Golden *et al.* 2017), we are not aware of other models focused specifically on targeting youth for antiretroviral treatment.

We hypothesized that youth-centered Treatment-as-Prevention (TasP) could yield large benefits—well in excess of that suggested by the models of age-based testing cited above. Our hypothesis is based on the following observations. (1) Young people are more likely than older people to have short-term relationships (Stignum *et al.* 1997, Johnson *et al.* 2001, Powers *et al.* 2011) (2) Young people who are in a partnership have higher coital frequency than older people (Call *et al.* 1995, Gray *et al.* 2001, Stewart *et al.* 2002, Brewis and Meyer, 2005). (3) Young people have behavioral and biological factors (such as higher rates of condom breakage and altered cytokine profiles at mucosal sites of entry) that raise the per-act probability of transmission (Goodkin *et al.* 2004, Valappil *et al.* 2005, Hughes *et al.* 2012, Yi *et al*. 2013). (4) Young people tend to partner with other young people (Barbieri and Hertrich 2005, Harling *et al.* 2014, and Harling *et al.* 2015). (5) People are rarely infected by HIV+ partners under the age of 16 (Shisana *et al.* 2014). These last two points suggest that treating people between 16 and 25 can result in “herd immunity” to teenagers entering the sexually active population (Bershteyn *et al.* 2016).

To test our hypothesis, we extended an existing agent-based HIV epidemic model for sexual networks to include the risk factors listed above (online technical supplement). To strike a balance between simulation speed and realism, we model a population of 2000 heterosexual individuals. Since key transmission rate parameters, such as the average number of partners per person, are independent of population size for sexually transmitted diseases (McCullum *et al.* 2001, Lloyd-Smith *et al.* 2004), our results should generalize to larger populations. The model implements a simplified treatment cascade: 100% of the reachable population (95% of the total population) is tested annually but only a subset (whose size we vary in our simulations) of diagnosed people are linked to effective care, resulting in a suppressed viral load. We assume that a proportion (default 5%) of treated individuals will drop out of care each year and no longer be virally suppressed (Yu *et al.* 2007, Fleishman *et al.* 2013, and Mberi *et al.* 2015). Dropouts who are a member of a target group have a higher chance of getting relinked to care than those who are not. In the absence of dropping out, however, individuals remain on care indefinitely even if they were originally linked to care due to a factor (e.g., youth) that no longer applies. Our default model assumes that all couples have the same probability of using a condom, an assumption that makes our default model most relevant (given uncertainties in model parameters) to a situation in which no one uses condoms. However, we also investigate variants of the model in which the probability of using a condom decreases with age, as reported in da Silveira *et al.* (2005), Simbayi *et al.* (2014), and Maticka-Tyndale E (2012).

Prior to the TasP campaign, a random subset of infected people receives suppressive therapy. The TasP campaign doubles the percentage of HIV+ people receiving suppressive therapy, *S,* from what it was 5 years before the TasP campaign (default 30%) to a TasP target, *S*_*targ*_ (default 60%). To simplify the accounting, we assume the TasP campaign is implemented instantly (rather than, for example, having six annual increases of ~12%). After the TasP target has been hit, the *absolute number* of individuals receiving suppressive therapy grows annually at rate *r* (default 2% per year) to account for population growth (default 1% per year) and general increases in economic productivity (default 1% per year). In simulations below, we focus on the success of TasP programs as a function of the targeting strategy and *S*_targ_. We note that *S*_*targ*_ gives the percentage of HIV+ people receiving suppressive therapy, *S*, at a single point in time (i.e., just after TasP target has been hit). During an unsuccessful TasP campaign (i.e., one that fails to reduce prevalence by year 30), the annual treatment growth term, *r*, can cause *S* to increase transiently before declining in the face of an expanding epidemic. For the simulations described in Figures 1 and 2 below, we found (data not shown) that, *S_max_*, the maximum percentage of HIV+ people receiving suppressive therapy following an unsuccessful TasP campaign exceeded *S_targ_* by an average of ~1% (e.g., *S_targ_* = 20% corresponding to *S_max_* ~21%).

We recognize that targeting individuals for treatment based on criteria other than their own health needs raises ethical concerns. However, HIV+ people sometimes fail to achieve viral suppression despite the best efforts of the existing health care system in a given locale (Kay *et al.*2016, Maman *et al.* 2016). Some patients do not return for care despite attempts by the clinic to contact them; i.e., are lost to follow-up (LTFU) (Wools-Kaloustian *et al*. 2006, de Almeida *et al.* 2014). Our model, in the first instance, could be viewed as deciding upon which of those LTFU should be prioritized for re-engagement with care. We note that many of those individuals LTFU (as well as many non-LTFU) patients require active interventions such as help with transportation, food-security, enrollment in support groups, adherence counseling (Fox *et al.* 2016), general HIV education (Katz and Bangsberg 2016), and/or other forms of health promotion (Psaros *et al.* 2015) before they can achieve viral suppression. Our model, therefore, can also encompass the effect of targeting promotion of such services within different subsets of the diagnosed, unsuppressed population (Table 1).

In Figure 1 we show the number of infected, the number treated, incidence, and AIDS death rates under the “CD4<500” and “under age 30” targeting strategies (as defined in Table 1), assuming *S_targ_* = 60% (i.e., that 60% of infected people receive suppressive therapy after starting the TasP campaign). In contrast to the “CD4<500” strategy where incidence increased over time, under the “under age 30” strategy incidence dropped to negligible levels (i.e., less than 0.1 infections per 100 person years) in 16/16 replicate simulations 23 years after the start of the TasP campaign. [Although viral loads drop to very low levels in “virally suppressed” people, they do not actually go to zero, meaning that it is not possible to stop all infections in the model even with 100% of HIV+ people being virally suppressed.] We note that the “CD4<500” strategy [as well as the “SPVL” (set point viral load) strategy in Figure 2], included for purposes of comparison, assumes assess to information that may not be available in resource-limited settings. For someone diagnosed as HIV+ who never returned for treatment, for example, CD4 count will probably not be known (though this could change with wider availability point-of-care CD4 tests).

**Table 1.**
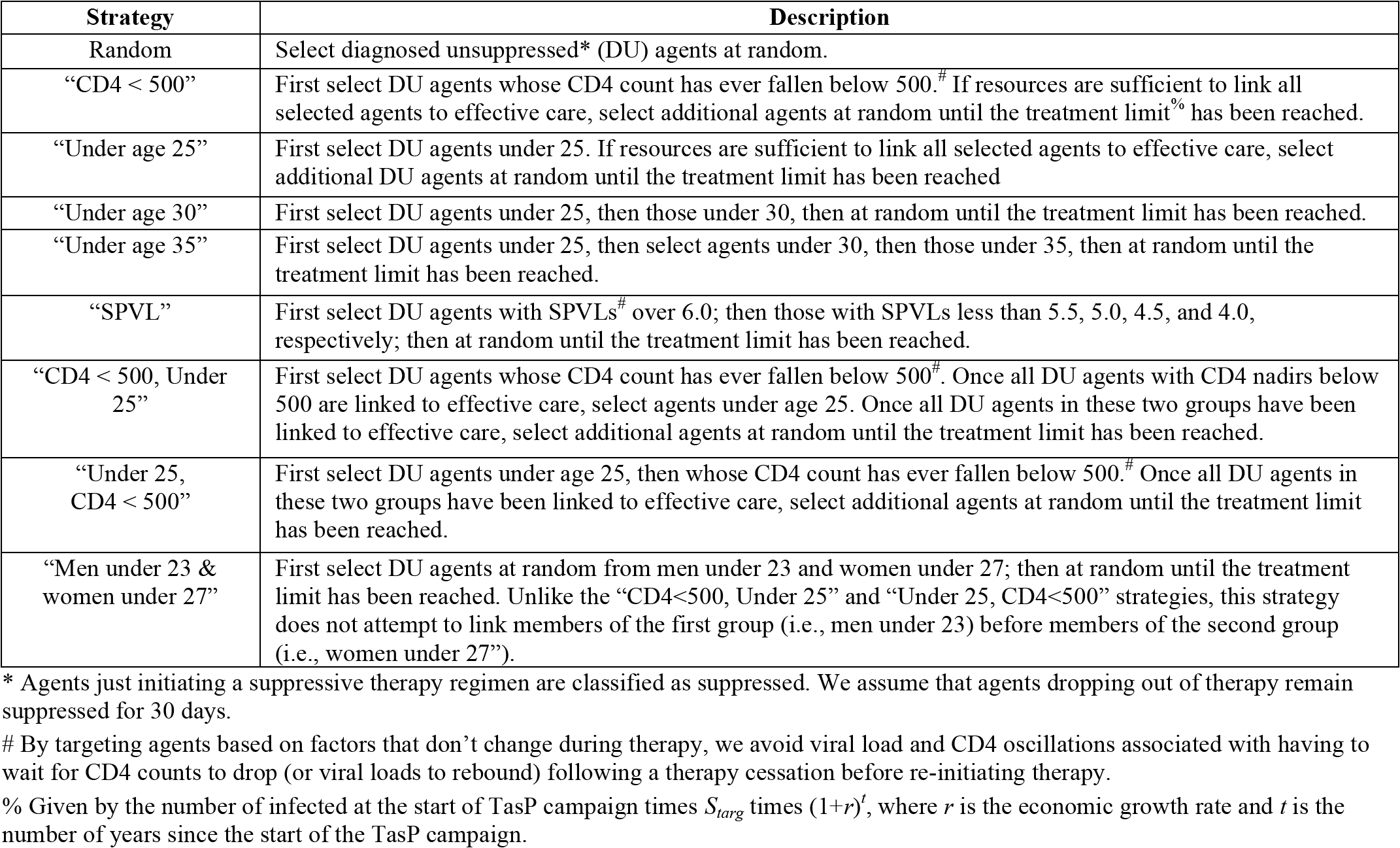
TasP Strategies modeled in this paper.

| Strategy | Description |
| --- | --- |
| Random | Select diagnosed unsuppressed* (DU) agents at random. |
| “CD4 < 500” | First select DU agents whose CD4 count has ever fallen below 500.<sup>#</sup> If resources are sufficient to link all selected agents to effective care, select additional agents at random until the treatment limit<sup>%</sup> has been reached. |
| “Under age 25” | First select DU agents under 25. If resources are sufficient to link all selected agents to effective care, select additional DU agents at random until the treatment limit has been reached |
| “Under age 30” | First select DU agents under 25, then those under 30, then at random until the treatment limit has been reached. |
| “Under age 35” | First select DU agents under 25, then select agents under 30, then those under 35, then at random until the treatment limit has been reached. |
| “SPVL” | First select DU agents with SPVLs<sup>#</sup> over 6.0; then those with SPVLs less than 5.5, 5.0, 4.5, and 4.0, respectively; then at random until the treatment limit has been reached. |
| “CD4 < 500, Under 25” | First select DU agents whose CD4 count has ever fallen below 500.<sup>#</sup> Once all DU agents with CD4 nadirs below 500 are linked to effective care, select agents under age 25. Once all DU agents in these two groups have been linked to effective care, select additional agents at random until the treatment limit has been reached. |
| “Under 25, CD4 < 500” | First select DU agents under age 25, then whose CD4 count has ever fallen below 500.<sup>#</sup> Once all DU agents in these two groups have been linked to effective care, select additional agents at random until the treatment limit has been reached. |
| “Men under 23 & women under 27” | First select DU agents at random from men under 23 and women under 27; then at random until the treatment limit has been reached. Unlike the “CD4<500, Under 25” and “Under 25, CD4<500” strategies, this strategy does not attempt to link members of the first group (i.e., men under 23) before members of the second group (i.e., women under 27”). |

\* Agents just initiating a suppressive therapy regimen are classified as suppressed. We assume that agents dropping out of therapy remain suppressed for 30 days.
<sup>#</sup> By targeting agents based on factors that don’t change during therapy, we avoid viral load and CD4 oscillations associated with having to wait for CD4 counts to drop (or viral loads to rebound) following a therapy cessation before re-initiating therapy.
<sup>%</sup> Given by the number of infected at the start of TasP campaign times $S_{targ}$ times $(1+r)^t$ , where $r$ is the economic growth rate and $t$ is the number of years since the start of the TasP campaign.

**Figure 1.**
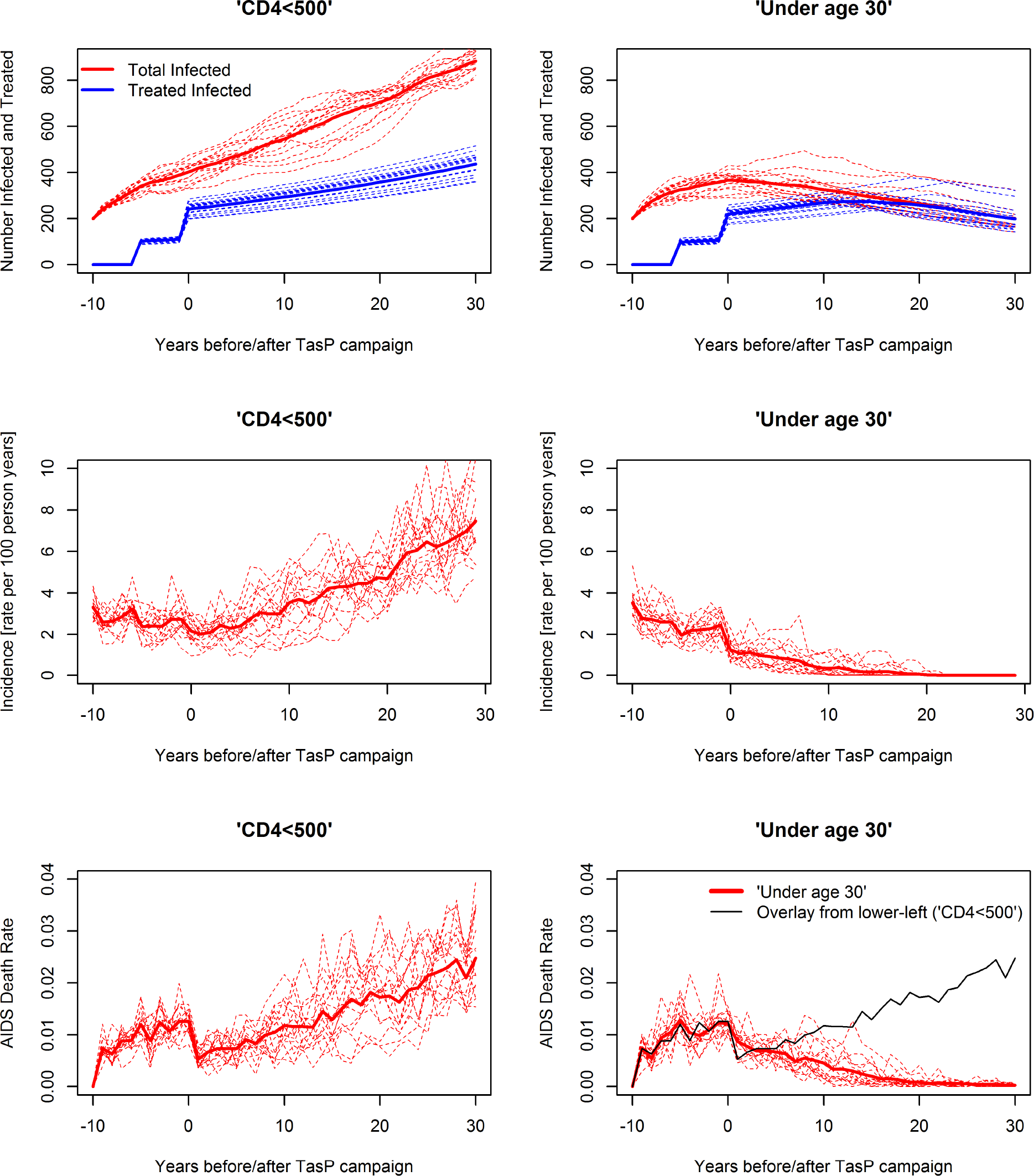
Simulations of the numbers of infected (red) and treated (blue) individuals (top panels), incidence (middle panels) and AIDS death rates (lower panels) following our “CD4<500” (left-side) and “under age 30” (right side) treatment-as-prevention (TasP) strategies. To account for existing treatment, we assume 30% of infected people (selected at random) receive suppressive therapy beginning five years before the TasP campaign. The TasP campaign immediately increases the percentage of HIV+ people receiving suppressive therapy to 60%. Once the TasP campaign starts, the model uses the CD4-or age-based strategies shown above to link a subset of unsuppressed diagnosed people to effective care. The 2% per year increase in the number treated after the TasP campaign was added to reflect population growth and increasing health care productivity over time. Thick lines give the mean of 16 independent replicates; thin dashed lines show the individual runs. The decline in the number treated after year 20 in the top right panel occurs because all “reachable infected agents” have been treated. The black line in the lower right panel shows for purposes of comparison the average number of AIDS deaths from the “CD4<500” strategy in the lower left panel.

In Figure 2a we show the average incidence 20-30 years after implementing the TasP campaign as a function of *S_targ_* for a broader range of strategies. For the “random” and “CD4<500” targeting strategies, we need to set *S_targ_* to 80% to reduce incidence rates to negligible levels (Figure 2a). Under the age-based targeting strategies, by contrast, incidence can be reduced to negligible levels using *S_targ_* values of 50% to 60% (Figure 2a). Comparable results were obtained using the total number of AIDS deaths over 30 years as a metric of impact (Figure 2b). The total number of person-years of treatment over the course of the simulation under age-based targeting strategies was comparable to the “SPVL” strategy but lower than it was under the “random” and “CD4<500” strategies (Figure 2c). This occurred because these strategies were more likely to reduce incidence to negligible levels, eliminating the necessity to treat additional people in later years.

Of the five “non-random” strategies in Figure 2, the percentage of the population initiating suppressive therapy at the start of the TasP campaign that was not a member of a target group was maximized under the “under age 25” strategy (Figure 2d). For *S_targ_* = 60%, for example, ~78% people who received suppressive therapy under “under age 25” strategy fell outside the target group, compared to ~41% under the “under age 30” strategy and only ~4% under the “SPVL” strategy (Figure 2d). If we include people who were virally suppressed prior to the “under age 25” TasP campaign, this percentage jumps to ~87%. The “under age 25” strategy, therefore, succeeds and has the potential to be a comparatively inclusionary strategy. In the subsequent simulations, we focus mainly on comparing the “under age 25” strategy with the “random” and/or “CD4<500” strategies assuming *S_targ_* = 60%.

The *y*-axis of Figure 2d is similar to *S_targ_* in that it quantifies “inclusivity” at a single point it time (i.e., just after the TasP target has been hit). For the TasP campaigns that reduced prevalence in Figure 2 (a subset that includes all of the simulations in which incidence dropped to zero), the percentage of untargeted individuals receiving suppressive therapy increased in subsequent years (data not shown). However, for the TasP campaigns that allowed prevalence to increase by more than the treatment growth rate *r*, the percentage of untargeted individuals receiving suppressive therapy decreased in subsequent years (data not shown). In other words, the successful TasP campaigns became more inclusionary over time, while unsuccessful TasP campaigns became more exclusionary over time. Fortunately, our simulations suggest that one will not have to wait long to find out whether a campaign will succeed. For the successful “under age 30” strategy shown in Figure 1, for example, we observed a ~2-fold drop in incidence within 2 years of starting the TasP campaign. While drops in incidence are unlikely be as rapid under more realistic (i.e., non-instantaneous) ramp-ups, incidence is a highly sensitive and general measure that can, with the aid of an epidemiological model, predict whether an in-progress TasP campaign is likely to reduce prevalence.

While age-based targeting greatly reduces the number of deaths over the long-term, it comes at the cost of allowing slightly more AIDS deaths relative to CD4-based targeting in the years immediately following the TasP campaign (Figure 1, lower-right panel, years 1-3; top and bottom rows of Table 2). The advisability of age-based targeting will depend, therefore, on whether public health officials take a long or short-term perspective. As shown in Figure 2, it also depends on the percentage of the population that can be virally suppressed. For high levels of suppression(i.e., *S_targ_* >=80%), the advantage to age-based targeting largely disappears. Age-based targeting, like most optimization schemes, becomes useful when resources are insufficient to control the epidemic using standard approaches.

**Figure 2.**
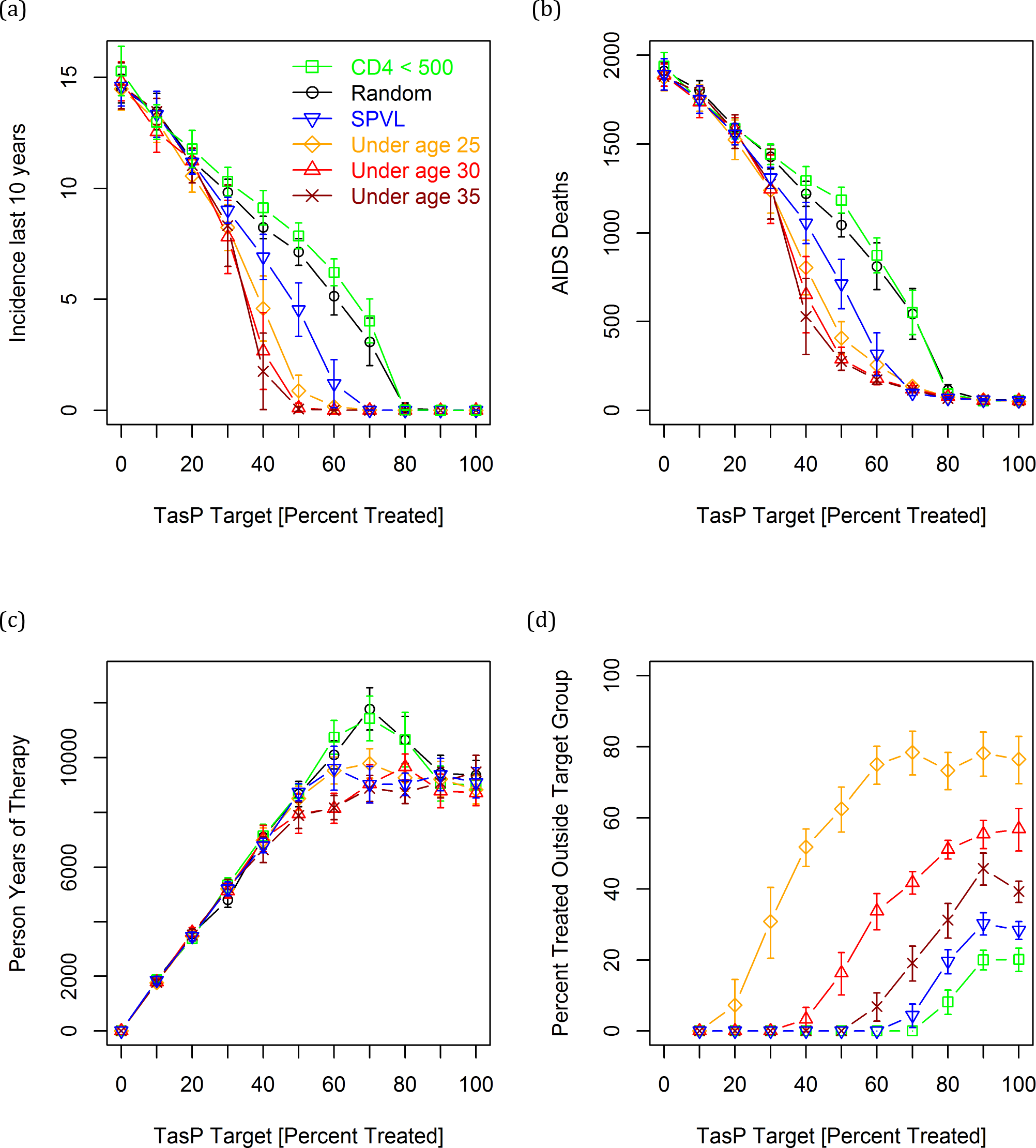
Simulations showing the effect of targeting strategy and *S_targ_* (percentage of infected people treated immediately after implementing the TasP campaign) on: (a) incidence 20-30 years after the TasP campaign, (b) AIDS deaths between years 0 and 30, (c) person-years of therapy (included to verify that the age-based strategies didn't somehow result in more people being treated), and (d) the percentage of people initiating suppressive therapy at the start of the TasP campaign who were not a member of a target group. Each point is the mean of 16 replicates. Error bars give approximate 95% confidence intervals. We note that the incidence in the absence of treatment in panel (a) matches peak levels in the heavily affected regions circa 1999 (Williams *et al.* 2001, Abdool Karim and Abdool Karim, 2002, Nel *et al.* 2012). Declines in the number of person years of therapy as *S_targ_* increased from 70 to 100% for “CD4<500” and “Random” reflect the ability of near universal therapy to reduce treatment expenditures over the long term.

**Table 2.** Tradeoff between short-and long-term outcomes involving age-and CD4-based strategies. Entries give mean and standard deviation of 16 replicates with *S_targ_* = 60%.

| Targeting Strategy | AIDS Deaths |  | New Infections |  |
| --- | --- | --- | --- | --- |
|  | Years 0-5 | Years 25-30 | Years 0-5 | Years 25-30 |
| CD4 < 500 | 85 (20) | 301 (56) | 212 (51) | 445 (44) |
| CD4 < 500, Under 25 | 81 (15)<sup>%</sup> | 286 (35) | 203 (26) | 436 (45)<sup>&</sup> |
| Under 25, CD4 < 500 | 87 (25)<sup>%</sup> | 25 (21) | 154 (36) | 49 (63) |
| Under age 25 | 104 (16) | 13 (14) | 148 (44) | 9 (18) |

<sup>%</sup> Significant reductions in AIDS deaths in years 0-5 compared to “Under age 25” ( $t$ -tests, $p < 0.001$ and $p < 0.05$ ).
<sup>&</sup> Significant increases in incidence in years 25-30 compared to “Under age 25” ( $t$ -tests, $p < 0.05$ ).

Our model includes many parameters, some of which are estimated with uncertainty. We therefore conducted a series of sensitivity analyses, focusing on parameters that we felt *a priori* might cause us to reject our hypothesis about the benefits of age-based TasP targeting. We find that age-based targeting continues to outperform random and CD4-based targeting in simulations in which: all agents have the same average relationship duration (Table 3), young people no longer have a higher per-act risk of infection (Table S1), young and old people have the same coital frequency (Figure S1), CD4 counts decline faster in older HIV+ patients in the absence of treatment (as noted, albeit cautiously, in Cori *et al.* 2015) (Figure S2), the frequency at which people are tested for HIV was varied (Figures 3 and S3), the rate at which untargeted agents drop out of care was varied (Figure S4), the average age difference between partners was varied (Figure S5), the percentage of people who could potentially be linked to care was varied (Figure S6), the probability of using a condom declines with age (Figure S7), and *r*, the annual increase in people receiving suppressive therapy after TasP target was hit, was varied between 0 and 9% per year (Figure S8). The one parameter that stood in all these simulations was testing frequency:although age-based targeting outperformed random and CD4-based targeting across a board range of testing frequencies, the key benefit to age-based targeting (i.e. the ability to drive incidence to zero in ~20 years with *S_targ_* = 60%) tapered off as testing intervals exceeded a year (Figure 3, Figure S3). Other than testing frequency, however, our results do not appear to be sensitive to any of our parameter values, at least when considered in isolation.

**Table 3.** Simulations in which all agents have the same average relationship duration (i.e., in a model in which young are no more likely than older people to have short relationships). Table entries show the mean and standard deviation in the number of infections during years 20 to 30 as a function of targeting strategy and the average relationship duration in a simulation with *S_targ_* = 60%.

| Targeting Strategy | Number of new infections in years 20-30<sup>#</sup> assuming an average relationship duration of |  |  |  |
| --- | --- | --- | --- | --- |
|  | 1 Years | 2 Years | 4 Years | 6 Years |
| CD4 < 500 | 743 (66) | 703 (103) | 749 (94) | 688 (170) |
| Random | 700 (101) | 703 (125) | 731 (64) | 663 (106) |
| Under age 25<sup>#</sup> | 3 (4) | 25 (82) | 22 (48) | 24 (66) |

<sup>#</sup> Significant reductions for “Under age 25” compared to “CD4<500” and “Random” for all six relationship durations ( $t$ -test, $p < 0.0001$ for all 8 comparisons). However, we see no differences by relationship duration for either “Under age 25”, “Random”, or “CD4<500” ( $t$ -tests, $p > 0.05$ all comparisons).

To test the model further, we performed a partial Latin hypercube analysis (Blower and Dowlatabadi 1994) to compare age-and CD4-based targeting strategies, assuming *S_targ_* = 60%, in simulations in which multiple parameters were randomized at the same time (Table S2). Age-based targeting reduced the total number of AIDS deaths in ~70% of these simulations (Figure 4). In the ~30% of cases where age-based targeting did not reduce the number of deaths relative to CD4-based targeting, differences were small and nonsignificant (see below for details). The relative performance of age-based targeting depended more on the strength of the epidemic than on any specific combination of age-related parameters. Our randomization process, being undirected, resulted in a number of either very weak or very strong epidemics. For very weak epidemics, defined as those that resulted in a final prevalence below 7%, there was a trend (*t*-test, *p* =0.14) for fewer AIDS deaths under CD4-based targeting; however, we can see from Figure 4 (bottom panel, left side) that any differences that might be shown to be significant under a larger simulation would be small. For very strong epidemics, defined as those that resulted in final prevalence greater than 70%, the advantage to age-base targeting, though technically significant according to a *t*-test (p=0.035), seemed go away. We note that a final prevalence of 70% is a catastrophic outcome that is unlikely to be reached in real life. These epidemics were so strong, in fact, that they could barely be controlled even after suppressing 100% of the reachable infected population (=95% of infected individuals) (data not shown). For simulations with more realistic final prevalences, we observed benefits to age-based targeting in 98% (92 out of 94) of simulations (Figure 4, bottom panel, middle section).

**Figure 3.**
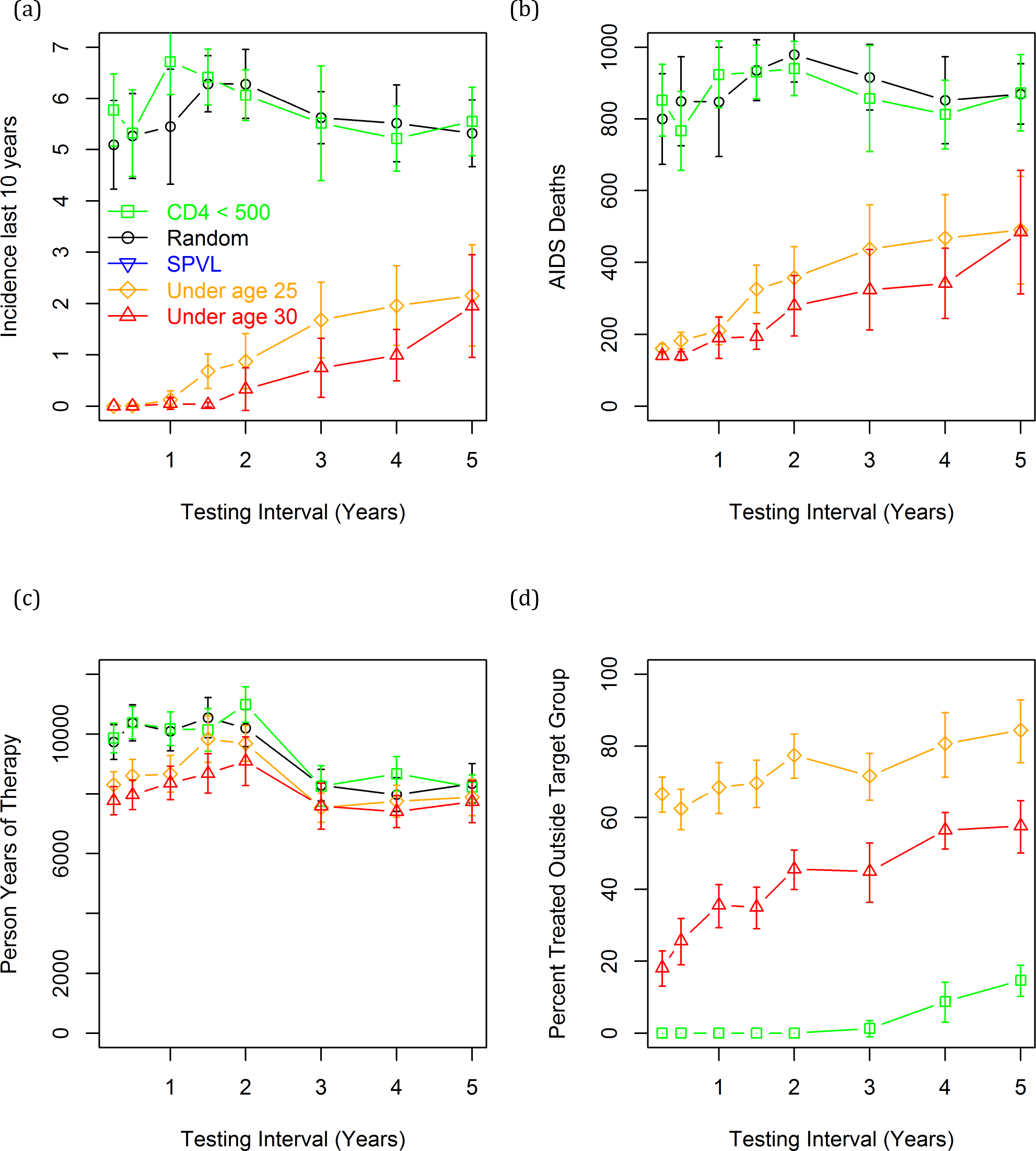
Effect of varying testing frequency on: (a) incidence 20-30 years after the TasP campaign, (b) AIDS deaths between years 0 and 30, (c) person-years of therapy, and (d) the percentage of people initiating suppressive therapy at the start of the TasP campaign who were not a member of a target group. This simulation assumes *S_targ_* = 60%. The thick lines give the mean of 16 replicates. Thin lines give mean plus and minus the standard error of the mean.

Our analyses have focused so far on strategies that consider age and CD4 status in isolation. Our model also allows us to model hybrid strategies. Compared to the “under age 25” strategy, we can reduce the number of short-term deaths by combining age and CD4 status (Table 2, middle rows); however, this comes at a cost of a slightly higher incidence in years 35-40. (The higher longterm incidence under the “Under 25, CD4 < 500” compared to “Under age 25” seems to be due to the model waiting to treat infected people until their CD4 counts has dropped, providing more time for an infected individual to transmit to his/her partner; however, we emphasize that this underperformance is slight and of borderline statistical significance). Use of hybrid strategies, in other words, failed to resolve the tradeoff illustrated in Figure 1 and Table 2 between short and long-term benefits in our model. We noted a trend towards better short and long-term outcomes relative to “CD4<500” using a strategy that first targeted those with CD4<500, then those under 25; however, the improvement in long-term outcomes associated with adding age to CD4 status was trivial compared to the effect of first targeting based on age. We conclude that targeting based on age is still the most effective long-term strategy.

**Figure. 4.**
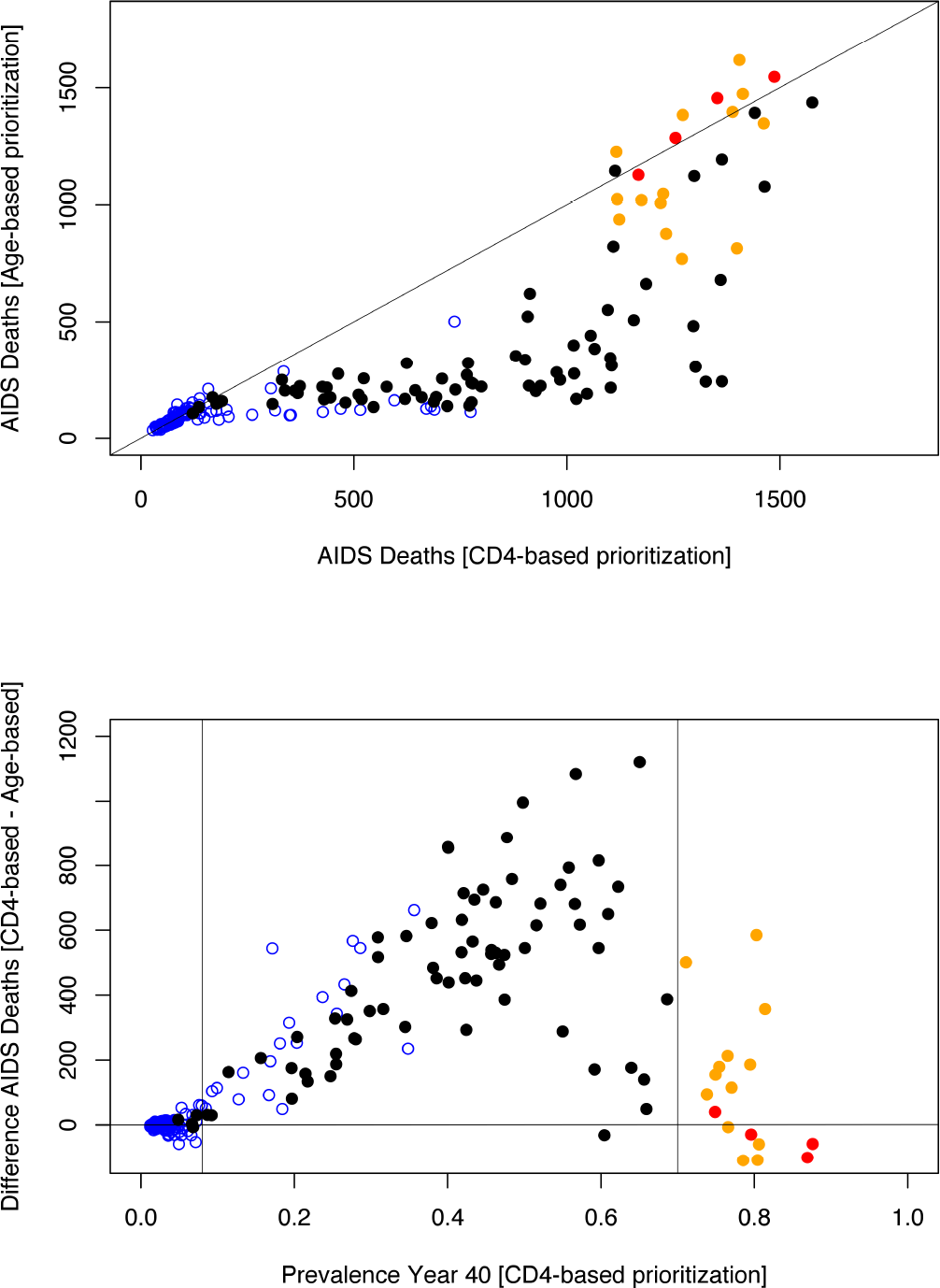
Number of AIDS deaths following age and CD4-based targeting strategies in simulations in which multiple parameters were randomized. Top: Total number AIDS deaths during years 0 to 30 under the “under age 25” strategy as a function of the total number AIDS deaths under the “CD4<500” strategy. Bottom: Difference between number of AIDS deaths following CD4-based targeting (“CD4<500”) and age-based targeting (“under age 25”) as a function of final (year 30) prevalence following CD4-based targeting. Each circle represents the end result of a set of simulations with a different set of parameter values. The blue points indicate simulations in which prevalence either decreased or increased by less than 50% prior to the TasP campaign. The orange and red points represent simulations in which prevalence increases >4-fold and >5-fold, respectively, prior to the TasP campaign. The thin vertical lines in the lower panel give year 30 prevalence ranges (7 and 70%) outside of which there was no longer a clear benefit to age-based targeting.

Our results come with the caveat that the failure to achieve 100% suppression is due primarily to steep drop-offs in steps in the treatment cascade that occur subsequent to testing. Under an alternative framework in which *testing* (rather than linkage to care) is the limiting factor, others have reported that targeting strategies may be optimally directed at slightly older people as prevalence tends to increase with age (Bershteyn *et al.* 2016, Golden *et al.* 2017). Another caveat is that our model ignores the tendency in many societies for younger women to partner with older men (Barbieri and Hertrich 2005, Harling *et al.* 2014). In such situations, it is reasonable to speculate that it might be advantageous to set a higher target age for men. In Figure S9, however, we show that altering the target ages for men and women has no discernable effect on either HIV incidence or AIDS deaths. While we leave open the possibility that a model that explicitly accounts for the tendency for women to pair with older men would show a benefit to targeting older men, this analysis suggests that any such benefit would be small relative to the large benefits that accrue to targeting young people.

In summary, our model suggests that under conditions where resources are projected to be inadequate to control the epidemic using existing targeting strategies (despite annual testing), that the addition of a new, supplemental program ensuring that people under 25 are linked to effective care, while maintaining (or preferably expanding) treatment rates for the general population could greatly enhance the success of treatment as prevention campaigns. Age-based targeting can, in principle, be facilitated by schools (Madiba and Mokgatle 2015), youth centers (Erulkar *et al.* 2001), military academies (Thomas *et al.* 2014), and universities (Huang 2016); however, there are many regions of the world where such institutions cover too low a percentage of the population (and/or lack the mandate or resources) to direct a sufficient percentage of young people to HIV testing and treatment services. In some of these regions, a demographic “bulge” associated with reductions in infant mortality since the year 2000 is stating to increase the number of young people at risk for acquiring HIV (UNICEF 2016, Avert 2017). Rigorous assessments of methods for targeting and engaging youth, retaining them in care, and keeping them virally suppressed are urgently needed, not just as an immediate humanitarian measure, but as a means for bringing the epidemic under the reproductive threshold and towards extinction.

## Methods

Our model builds on “Evonet-HIV”, an agent-based simulation model for HIV epidemiology originally developed for the purpose of modeling SPVL evolution (Herbeck *et al.* 2017, Goodreau *et al.* 2017, Stansfield *et al.* 2017). Each “agent” has a series of attributes including age, sex, HIV status, viral load, CD4 count (discretized into 4 bins), and HIV diagnosis status. Evonet-HIV includes generalizable ERGM (exponential random graph model) terms for the formation and dissolution of sexual partnerships, and includes modules for risk group behavior, HIV testing, viral load change, the effect of viral load on CD4 decline and HIV transmission, and the effect of treatment on viral load. The code can be accessed at https://github.com/EvoNetHIV/Mittler-et-al-TasP-by-Age.

Detailed methods, data sources, and methods for fitting the model to data are given in the <u>online-supplemental-methods</u>. Briefly, we assume an initial population of 2000 people of whom an average of 200 are HIV infected (i.e., 10% prevalence). HIV-negative people enter the sexually active population at 16 and die of natural causes according to age-specific mortality tables in the absence of HIV. In the absence of HIV-induced mortality the population increases each year (default 1% per year). With our default parameters, the number of infected people doubles every ~10 years in the absence of treatment.

To mimic the tendency of young people to have short-term relationships, we create an attribute for risk group membership (1 = no steady partner, occasional short-term relationships; 2 = tendency to form moderately long-term partnerships; 3 = tendency to form long-term partnerships). 90% of agents entering the population at age 16 belong to groups 1 and 2. Agents in risk groups 1 and 2 are assumed to have small per-day probabilities of transitioning to risk groups 2 and 3; i.e., agents have a tendency to have longer partnerships as they age. To mimic the tendency of young people to partner with other young people, we added an ERGM term that forced the average age difference in the population to equal a specified value (default 4 years). Higher probabilities of sex and transmission in younger people were modeled using straightforward modifications to our modules for the probability of sex and transmission.

Into this mix, we introduce a TasP campaign that targets different kinds of individuals for linkage to effective care. The model assumes an overall treatment limit given by *S_targ_I*_0_(1 + *r*)^*t*^, where *S_targ_* is the percent of infected people linked to effective care at the start of the TasP campaign, *I_0_* is the number of infected people at the start of the TasP campaign, *r* is the economic growth rate, and *t*=the number of years since the start of the TasP campaign. Targeting strategies include: age (e.g. “under age 25”), immunological status (e.g., “CD4<500”), combinations of age and immunological status (e.g., “Under 25, CD4 < 500”), viral load (e.g., “SPVL”), and no factor at all (“random”) (Table 1). Several of the strategies include a targeting hierarchy within the primary target range (Table 1). “Under age 30”, for example, targets those under age 25 before those between ages 25 and 30. In these cases, agents are linked to care at random within each successive target group until the overall treatment limit is reached. To ensure that equal numbers are treated under all strategies (prior to one strategy linking 100% of people to effective care), all strategies include a final “random” component that is applied once all of the people in the primary target groups have been linked to effective care.

## Acknowledgments

We thank Neil Abernethy for comments. This work was supported by the National Institutes of Health, through grants R01-AI108490, R21-HD075662, and R01-HD068395. This work was facilitated though the use of advanced computational, storage, and networking infrastructure provided by the Hyak supercomputer system and funded by the Student Technology Fee at the University of Washington. Partial support for this research came from an NICHD research infrastructure grant (R24-HD042828) to the Center for Studies in Demography and Ecology at the University of Washington. The content is solely the responsibility of the authors and does not necessarily represent the official views of the National Institutes of Health.

## References

Abdool Karim Q, Abdool Karim SS (2002) The evolving HIV epidemic in South Africa. International Journal of Epidemiology 31:37–40.

Avert 2017. Young people, HIV and AIDS. https://www.avert.org/professionals/hiv-social-issues/key-affected-populations/young-people (downloaded October 17, 2017)

Barbieri M and Hertrich V. (2005) Age Difference between Spouses and Contraceptive Practice in Sub-Saharan Africa. Population 60:617–654.

Bershteyn A, Klein, DJ, and Eckhoff, PA. (2016) Age-targeted HIV treatment and primary prevention as a ‘ring fence’ to efficiently interrupt the age patterns of transmission in generalized epidemic settings in South Africa. International Health 8:277–285.

Blower SM, Dowlatabadi (1994) Sensitivity and uncertainty analysis of complex models of disease transmission: an HIV model, as an example. International Statistical Review 62:229–243.

Brewis A, Meyer M (2004) Martial coitus across the life course. J. Biosoc Sci 37:499–518.

Call V, Sprech S, and Schwartz P. (1995) The Incidence and Frequency of Marital Sex in a National Sample Journal of Marriage and Family 57: 639–652.

da Silveira MF, dos Santos IS, Béria JU, Horta BL, Tomasi E, and & Victora CG (2005). Factors associated with condom use in women from an urban area in southern Brazil. Cadernos de Saúde Pública, 21:1557–1564.

de Almeida MC, de Jesus Pedroso N, do Socorro Lina van Keulen M, Jácome GP, Fernandes GC, Yokoo EM, Tuboi SH. (2014) Loss to follow-up in a cohort of HIV-infected patients in a regional referral outpatient clinic in Brazil. AIDS Behav. 18:2387–2396.

Erulkar AS, Beksinska M, Cebekhulu A (2001) An Assessment of Youth Centres in South Africa https://www.iywg.org/sites/iywg/files/SouthAfrica_Youth_Centers.pdf

Fleishman JA, Yehia BR, Moore RD, Korthuis PT, Gebo KA. (2012) Establishment, retention, and loss to follow-up in outpatient HIV care. J Acquir Immune Defic Syndr. 60:249–259.

Fox MP, Berhanu R, Steegen K, Firnhaber C, Ive P, Spencer D, Mashamaite S, Sheik S, Jonker I, Howell P, Long L, Evans D. (2016) Intensive adherence counselling for HIV-infected individuals failing second-line antiretroviral therapy in Johannesburg, South Africa. Trop Med Int Health. 21:1131–1137.

Golden MR, Hughes JP, Dombrowski JC. (2017) Optimizing the timing of HIV screening as part of routine medical care. AIDS Patient Care STDS. 31:27–32.

Goodkin K, Shapshak P, Asthana D, Zheng W, Concha M, Wilkie FL, Molina R, Lee D, Suarez P, Symes S, Khamis I. (2004) Older age and plasma viral load in HIV-1 infection. AIDS 18 Suppl 1:S87–98.

Goodreau SM, Stansfield SE, Murphy JT, Peebles KC, Gottlieb GS, Abernethy NF, Herbeck JT, Mittler JE. (2017) Sexual network structure, HIV prevalence, and the evolution of set point viral load. Submitted.

Gray RH, Wawer MJ, Brookmeyer R, Sewankambo NK, Serwadda D, Wabwire-Mangen F, Lutalo T, Li X, vanCott T, Quinn TC, and the Rakai Project Team (2001) Probability of HIV-1 transmission per coital act in monogamous, heterosexual, HIV-1-discordant couples in Rakai, Uganda. Lancet 357:1149–53.

Harling G, Newell ML, Tanser F, Kawachi I, Subramanian SV, Bärnighausen T. (2014) Do age-disparate relationships drive HIV incidence in young women? Evidence from a population cohort in rural KwaZulu-Natal, South Africa. J Acquir Immune Defic Syndr. 66:443–451.

Harling G, Newell ML, Tanser F, Bärnighausen T. (2015) Partner age-disparity and HIV incidence risk for older women in rural South Africa. AIDS Behav. 19:1317–1326.

Herbeck JT, Peebles KC, Edlefsen P, Rolland M, Stansfield SE, Murphy JT, Gottlieb GS, Abernethy NF, Mullins JI, Mittler JE, and Goodreau SM. (2017) HIV population-level adaptation can rapidly diminish the impact of a partially effective vaccine. bioRxiv110793.

Hontelez JA, Lurie MN, Bärnighausen T, Bakker R, Baltussen R, Tanser F, Hallett TB, Newell ML, de Vlas SJ. (2013) Elimination of HIV in South Africa through expanded access to antiretroviral therapy: a model comparison study. PLoS Med. 10, e1001534.

Huang E (2016) HIV is growing so fast among chinese youth that a university is selling testing kits in vending machines. https://qz.com/817266/hiv-is-growing-so-fast-among-chinese-youth-that-a-university-is-selling-testing-kits-in-vending-machines/

Hughes JP, Baeten JM, Lingappa JR, Magaret AS, Wald A, de Bruyn G, Kiarie J, Inambao M, Kilembe W, Farquhar C, Celum C. (2012) Partners in Prevention HSV/HIV Transmission Study Team. Determinants of per-coital-act HIV-1 infectivity among African HIV-1-serodiscordant couples. J InfectDis. 205:358–365.

Johnson AM, Mercer CH, Erens B, Copas AJ, McManus S, Wellings K, Fenton KA, Korovessis C, Macdowall W, Nanchahal K, Purdon S, Field J. (2001) Sexual behaviour in Britain: partnerships, practices, and HIV risk behaviours. Lancet 358:1835–1842.

Katz IT and Bangsberg DR (2016) Cascade of Refusal—What does it mean for the future of treatment as prevention in sub-Saharan Africa? Curr HIV/AIDS Rep. 13:125–130.

Kay ES, Batey DS, Mugavero MJ (2016) The HIV treatment cascade and care continuum: updates, goals, and recommendations for the future. AIDS Research and Therapy 13:35.

Kretzschmar ME, Schim van der Loeff MF, Birrell PJ, De Angelis D, Coutinho RA. (2013) Prospects of elimination of HIV with test-and-treat strategy. Proc Natl Acad Sci USA. 2013 110:15538–15543

Lloyd-Smith JO, Getz WM, Westerhoff HV. (2004) Frequency-dependent incidence in models of sexually transmitted diseases: portrayal of pair-based transmission and effects of illness on contact behaviour. Proc. R. Soc. Lond. B 271:625–635.

Madiba S, Mokgatle M. (2015) “Students want HIV testing in schools” a formative evaluation of the acceptability of HIV testing and counselling at schools in Gauteng and North West provinces in South Africa. BMC Public Health. 15:388.

Maman D, Chilima B, Masiku C, Ayouba A, Masson S, Szumilin E, Peeters M, Ford N, Heinzelmann A, Riche B, Etard JF. (2016) Closer to 90-90-90. The cascade of care after 10 years of ART scale-up in rural Malawi: a population study. J Int AIDS Soc 19:20673.

Maticka-Tyndale E (2012) Condoms in sub-Saharan Africa. Sexual Health 9:59–72.

Mberi MN, Kuonza LR, Dube NM, Nattey C, Manda S, Summers R. (2015) Determinants of loss to follow-up in patients on antiretroviral treatment, South Africa, 2004-2012: a cohort study. BMC Health Serv Res. 15:259.

McCallum H, Barlow N, Hone J. (2001) How should pathogen transmission be modelled? Trends in Ecology & Evolution 16:295–300.

Medlock J, Pandey A, Parpia AS, Tang A, Skrip LA, Galvani AP. (2017) Effectiveness of UNAIDS targets and HIV vaccination across 127 countries. Proc Natl Acad Sci USA. 114:4017–4022.

Nel A, Mabude Z, Smit J, Kotze P, Arbuckle D, Wu J., van Niekerk N, van Wijgert J. (2012) HIV incidence remains high in KwaZulu-Natal, South Africa: Evidence from three districts. PLoS ONE 7, e35278.

Powers KA, Hoffman IF, Ghani AC, Hosseinipour MC, Pilcher CD, Price MA, Pettifor AE, Chilongozi DA, Martinson FE, Cohen MS, Miller WC (2011) Sexual partnership patterns in Malawi: Implications for HIV/STI transmission. Sexually Transmitted Diseases 38:657–66

Psaros C, Remmert JE, Bangsberg DR, Safren SA, Smit JA. (2015) Adherence to HIV care after pregnancy among women in sub-Saharan Africa: falling off the cliff of the treatment cascade. Curr HIV/AIDS Rep 12:1–5.

Rozhnova G, van der Loeff MF, Heijne JC, Kretzschmar ME (2016) Impact of heterogeneity in sexual behavior on effectiveness in reducing HIV transmission with test-and-treat strategy. PLoS Comput Biol. 12, e1005012.

Shisana, O, Rehle, T, Simbayi LC, Zuma, K, Jooste, S, Zungu N, Labadarios, D, Onoya, D et al. (2014) South African National HIV Prevalence, Incidence and Behaviour Survey, 2012. Cape Town, HSRC Press.

Simbayi LC, Matseke G, Wabiri N, Ncitakalo N, Banyini M, Tabane C, Tshebetshebe D. (2014) Covariates of condom use in South Africa: findings from a national population-based survey in 2008. AIDS Care. 26:1263–1269.

Stansfield SE, Mittler JE, Gottlieb GS, Murphy JT, Hamilton DT, Detels R, Wolinsky SM, Jacobson LP, Margolick JB, Rinaldo CR, Herbeck JT, Goodreau SM. (2017) Sexual role and HIV-1 set point viral load among men who have sex with men. Submitted.

Stewart H, Morison L, White R (2002) Determinants of coital frequency among married women in Central African Republic: The role of female genital cutting. J. Biosoc Sci 34: 525–539.

Stignum H, Margnus P, Harns JR, Samuelsen SO, and LS Bakketeig (1997) Frequency of sexual partner change in a Norwegian population. American Journal of Epidemiology 145:636–643.

Thomas AG, Grillo MP, Djibo DA, Hale B, Shaffer RA. (2014) Military HIV policy assessment in sub-Saharan Africa. Mil Med. 179:773–777.

UNAIDS (2017) Ending AIDS: Progress towards the 90-90-90 targets. Downloaded from http://www.unaids.org/sites/default/files/mediaasset/GlobalAIDSupdate2017en.pdf on Oct 6, 2017.

UNICEF (2016) For Every Child, End AIDS-Seventh Stocktaking Report, United Nations Children's Fund, New York, December 2016.

Valappil T, Kelaghan J, Macaluso M, Artz L, Austin H, Fleenor ME, Robey L, Hook EW 3rd. (2005) Female condom and male condom failure among women at high risk of sexually transmitted diseases. Sex Transm Dis. 32:35–43.

Williams B, Gouws E, Wilkinson D, Abdool Karim SS. (2001) Estimating HIV incidence rates from age prevalence data in epidemic situation. Stat Med 20: 2003–2016.

Williams BG and Gouws E (2013) R0 and the elimination of HIV in Africa: Will 90-90-90 be sufficient? arXiv:1304.3720.

Wools-Kaloustian K, Kimaiyo S, Diero L, Siika A, Sidle J, Yiannoutsos CT, et al. (2006) Viability and effectiveness of large-scale HIV treatment initiatives in sub-Saharan Africa: Experience from western Kenya. AIDS 20:41–48.

Yi TJ, Shannon B, Prodger J, McKinnon L, Kaul R. (2013) Genital immunology and HIV susceptibility in young women. Am J Reprod Immunol 69 (Suppl. 1): 74–79.

Yu JK, Chen SC, Wang KY, et al. 2007. True outcomes for patients on antiretroviral therapy who are “lost to follow-up” in Malawi. Bull. World Health Organ. 85: 550–554.

